# Sleep rescues age-associated loss of glial engulfment

**DOI:** 10.1101/2025.04.02.646667

**Authors:** Jiwei Zhang, Elizabeth B. Brown, Evan Lloyd, Isabella Farhy-Tselnicker, Alex C. Keene

## Abstract

Neuronal injury due to trauma or neurodegeneration is a common feature of aging. The clearance of damaged neurons by glia is thought to be critical for maintenance of proper brain function. Sleep loss has been shown to inhibit the motility and function of glia that clear damaged axons while enhancement of sleep promotes clearance of damaged axons. Despite the potential role of glia in maintenance of brain function and protection against neurodegenerative disease, surprisingly little is known about how sleep loss impacts glial function in aged animals. Axotomy of the *Drosophila* antennae triggers Wallerian degeneration, where specialized olfactory ensheathing glia engulf damaged neurites. This glial response provides a robust model system to investigate the molecular basis for glial engulfment and neuron-glia communication. Glial engulfment is impaired in aged and sleep-deprived animals, raising the possibility that age-related sleep loss underlies deficits in glial function. To define the relationship between sleep- and age-dependent reductions in glial function, we restored sleep to aged animals and examined the effects on glial clearance of damaged axons. Both pharmacological and genetic induction of sleep restores clearance of damaged neurons in aged flies. Further analysis revealed that sleep restored post-injury induction of the engulfment protein Draper to aged flies, fortifying the notion that loss of sleep contributes to reduced glial-mediated debris clearance in aged animals. To identify age-related changes in the transcriptional response to neuronal injury, we used single-nucleus RNA-seq of the central brains from axotomized young and old flies. We identified broad transcriptional changes within the ensheathing glia of young flies, and the loss of transcriptional induction of autophagy-associated genes. We also identify age-dependent loss of transcriptional induction of 18 transcripts encoding for small and large ribosomal protein subunits following injury in old flies, suggesting dysregulation of ribosomal biogenesis contributes to loss of glial function. Together, these findings demonstrate a functional link between sleep loss, aging and Wallerian degeneration.

## Introduction

Sleep is a universal behavior that is critical for diverse aspects of brain function [1]. Chronic sleep disturbances are associated with numerous health consequences including neurodegenerative disease and cognitive decline [2,3]. Neurite damage due to apoptosis, trauma, or genetic factors is a common feature of aging, and the clearance of damaged neurons is thought to be critical for maintenance of proper brain function [4,5]. In the central and peripheral nervous systems, damaged neurites are cleared by Wallerian degeneration, a process where microglia or macrophages and Schwan cells are activated and engulf damaged neurites [6]. Despite the critical role of neurite clearance in maintenance of brain function and protection against neurodegenerative disease, surprisingly little is known about how life-history traits and environment modulate neurite clearance [7].

In flies and mammals, axotomy triggers Wallerian degeneration, where specialized glia engulf damaged neurites [8]. The genetic accessibility of the *Drosophila* olfactory system, combined with powerful genetic tools that allow for cell-type specific manipulation of gene expression, have made *Drosophila* a leading model to study injury-induced glial engulfment [9]. In *Drosophila,* olfactory ensheathing glia surround the antennal lobe then engulf the damaged olfactory neurons. Genetic screens in this system have identified numerous novel genetic factors and intercellular signaling pathways required for glial engulfment including *Stat92E, Draper, and Insulin Receptor* [10–14]. Further, numerous factors have been identified that function within neurons to signal neural injury including loss of the NAD+ synthase *Nmat1* and d*Sarm1* [9,12,15]. Modeling of Wallerian degeneration in other systems including neurons in the *Drosophila* wing, larval peripheral neurons, and mouse peripheral nerves confirm the mechanisms regulating Wallerian degeneration in the olfactory system are conserved in other *Drosophila* and mammalian neurons [16–18].

Glia play a critical role in the regulation of sleep duration, sleep-mediated neuronal homeostasis, and clearance of toxic substances during sleep [16–20]. Multiple mechanisms link glia to neurodegenerative disease including a role for clearance of Amyloid β (Aβ) by the glymphatic system and pruning of synapses by microglia in Alzheimer’s disease [21–23]. Recent findings suggest microglia function is inhibited by sleep deprivation in mouse models, raising the possibility that sleep directly impacts neuronal pruning and clearance within the brain [24]. Despite these bidirectional interactions between glia and sleep, surprisingly little is known about the mechanisms through which sleep deprivation impacts glial function, and how this contributes to neurodegenerative disease.

Growing evidence suggests sleep loss contributes to many of the functional deficits associated with aging [25,26]. Sleep loss shortens lifespan and accelerates many factors associated with aging, while quality and duration are reduced in old individuals [26,27]. In *Drosophila* aging, high calorie diets that accelerate aging, as well as acute sleep loss, all inhibit Draper expression, resulting in reduced glial plasticity and clearance of injured neurites [28–31]. While the effects of age and sleep-related factors have largely been studied independently, both result in disrupted sleep, raising the possibility that reduced sleep quality contributes to age-related decline in glial function [32–34]. Examining the role of sleep on aging-associated loss of glial engulfment has potential to guide experimental approaches and therapeutic treatments for age-related decline in brain function.

Here, we examine the relationship between sleep and age-dependent changes on glial engulfment of damaged neurites. Both pharmacological and genetic induction of sleep restore glial plasticity and Draper induction in aged flies. Further, the sleep-inducing effects of neural injury are absent in aged flies, revealing bidirectional interactions between sleep and aging. Together, these findings suggest sleep loss plays a critical role in age-dependent loss of glial cell function.

## Results

Sleep duration and quality are reduced in aged flies [27,34–37]. To identify the specific timepoints when sleep duration and quality are reduced, we compared sleep in flies aged 5, 25 and 40 days. There was a significant reduction in sleep in flies aged 25 days, and a greater loss at 40 days (Fig 1A,B). These changes in sleep duration are associated with reduced bout length and increased bout number, indicating reduced sleep quality (Fig S1A-F). Therefore, these timepoints were used to investigate the effects of age-related sleep loss on Wallerian degeneration.

**Figure 1.**
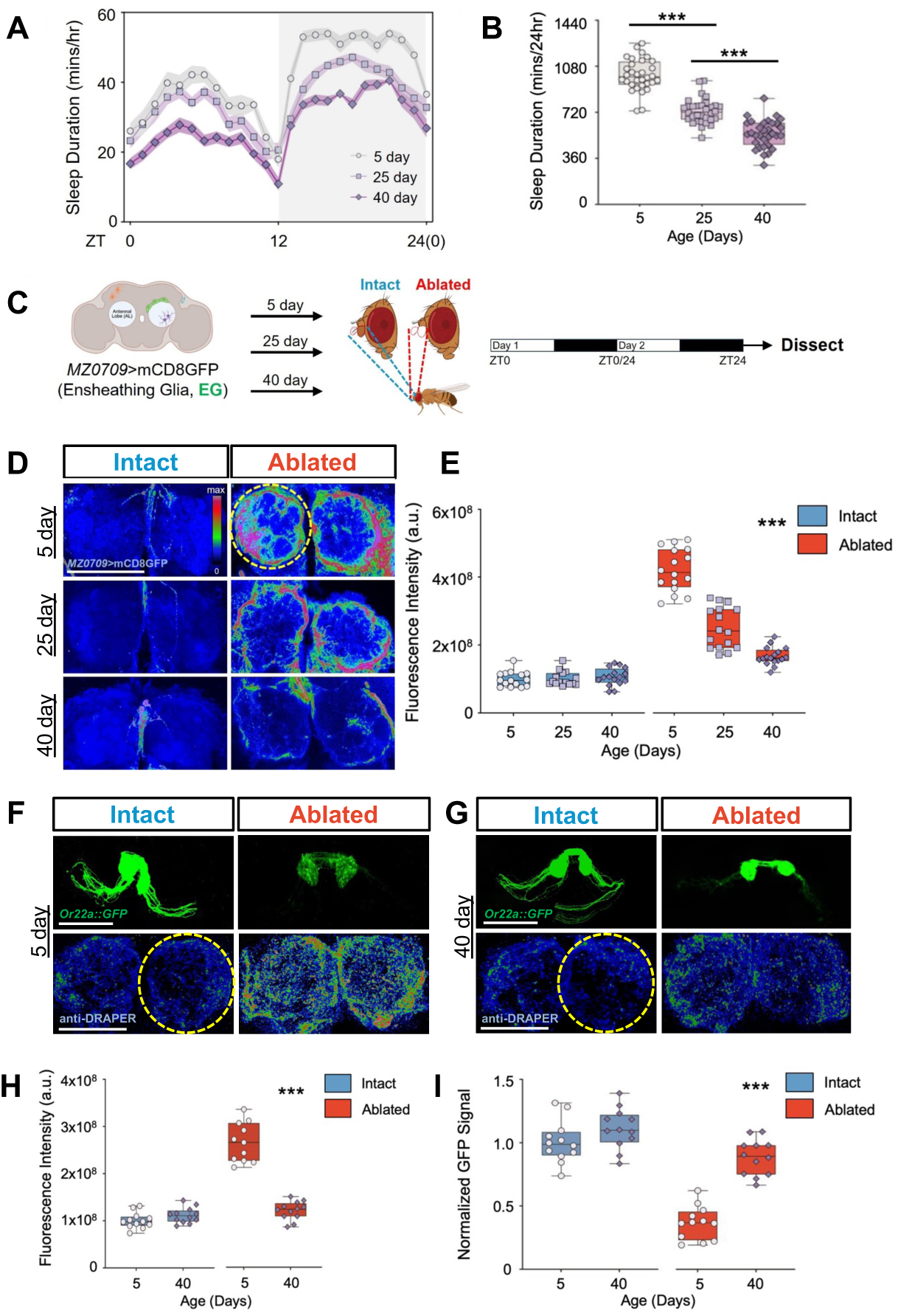
Glial innervation to neuronal injury is reduced in the aged flies. (**A**,**B**) Sleep profiles and sleep duration of flies at 5 days (white, cycle), 25 days (mauve, square), and 40 days of age (purple, diamond), n=32-36. (**C**) Schematic of experimental design for ensheathing glia (EG) response to neuronal injury in young (5-day-old) and aged (25-day-old or 40-day-old) flies. (**D**) Ensheathing glial innervation following antennal ablation in the antennal lobes of *MZ0709*-GAL4>mCD8GFP flies at 5, 25 and 40 days of age. Red circle indicates the region of the ensheathing glial membrane surrounding the antennal lobe that was quantified. (**E**) Quantification of glial innervation reveals a significant increase in ablated flies (red box) compared to unablated controls (blue box, two-way ANOVA, F_(2,84)_=77.69, P<0.001). There was an age-dependent decline in fluorescence intensity at 25-days (P<0.001) and 40-days of age (P<0.001), compared to 5-day axotomized flies. (**F-I**) GFP signal labeled by Or22a::GFP and (**F’-I’**) immunostaining for DRAPER in the antennal lobes of intact and ablated flies at 5- and 40-days of age. Hashed red circle represents the area of DRAPER quantification surrounding the antennal lobe and all fluorescence intensity were normalized to the 5-day intact controls. (**J**) DRAPER expression was significantly elevated in ablated 5-day-old flies compared to all other groups (P<0.001). (**K**) There was a significant reduction in GFP intensity in ablated 5-day old flies compared to all other groups (P<0.001, two-way ANOVA, F_(1,44)_=21.06). Tukey’s multiple comparison tests: *P<0.05; **P<0.01; ***P<0.001. Error bars indicate ± SEM. Scale bar denotes 50 μm.

Previous findings suggest that glial plasticity and Draper induction are impaired in aged flies [29]. To further examine the effects of aging on glial function we quantified Draper induction, neurite clearance and glial membrane plasticity in young and old flies. We ablated the antennae of flies aged 5, 25, and 40 days, expressing mCD8:GFP under control of the pan-glial driver Repo (Fig S1G) [9]. Glial membranes labeled with GFP robustly innervate the antennal lobes 24 hours following antennal ablation in young 5-day-old flies (Fig S1H), while membrane innervation of the antennal lobes is diminished in flies injured at 25 days old, with an additional reduction in 40-day-old injured flies (Fig S1I). To confirm the phenotypes observed were due to motility of olfactory ensheathing glia, we selectively labeled ensheathing glia with MZ0709-GAL4 and quantified antennal lobe innervation following antennal ablation (Fig 1C), [38]. Similar to results with Repo-GAL4, antennal innervation following injury was reduced at 25 days, and reduced further in ablated 40-day-old flies, while there was no age-dependent differences in intact flies (Fig 1D,E). These findings confirm that ensheathing glial innervation of the antennal lobe is impaired in aged flies.

Neuronal injury results in upregulation of the engulfment protein Draper within olfactory ensheathing glia, leading to clearance of the damaged neurites [9]. To confirm that age-related deficits in glial plasticity is associated with loss of Draper and reduced neurite engulfment, we genetically labeled the olfactory receptor neurons (ORNs) using an Or22a::GFP transgene that exclusively labels olfactory receptor neurons (ORNs) from the antennae, but not the maxillary palp [39]. The antennae of 5- or 40-day-old flies were ablated and the brains were subsequently immunostained for Draper 24 hours following injury. There were no differences in Draper levels in 5- and 40-day-old intact flies (Fig 1F-I, J). Antennal ablation robustly induced Draper within the antennal lobes of 5-day-old flies, but not 40-day-old flies, confirming Draper induction is reduced in aged flies (Fig 1F’-I’, J). Conversely, in 5-day-old flies, there was a significant reduction in Or22a::GFP signal, indicating robust glial engulfment (Fig 1F,G,K). Or22a::GFP signal was significantly higher in ablated 40-day-old flies compared to 5-day-old flies, revealing reduced glial engulfment of damaged neurites in aged flies (Fig 1H,I,K). Together, these findings confirm that glial motility, Draper induction, and neurite engulfment in response to injury are reduced in 40-day-old flies.

Injury and immune activation are associated with increased sleep in *Drosophila* [18,34]. In flies, sleep is increased immediately following antennal ablation, providing a system to examine the effects of aging on neural injury response [18]. To examine the effects of aging on sleep following injury, we quantified sleep immediately following injury in 5- and 40-day-old flies (Fig 2A). In agreement with previous findings, 5-day-old antennal ablated flies slept 182 significantly more than intact controls (Fig 2B) [18]. Conversely, there was no increase in daytime sleep following ablation in 40-day-old ablated flies (Fig 2C,D, S2A). In 40-day-old flies there was an increase in nighttime sleep 12 hours from injury, suggesting a reduced and delayed response (Fig S2A). At day two following ablation sleep did not differ from controls for both young and old flies, revealing the immediate effects of injury do not result in long-lasting changes (Fig 2B,C). The increased sleep following 188 injury in 5-day-old flies is due to longer bout lengths, consistent with greater sleep consolidation (Fig 2D, S2D-K).

**Figure 2.**
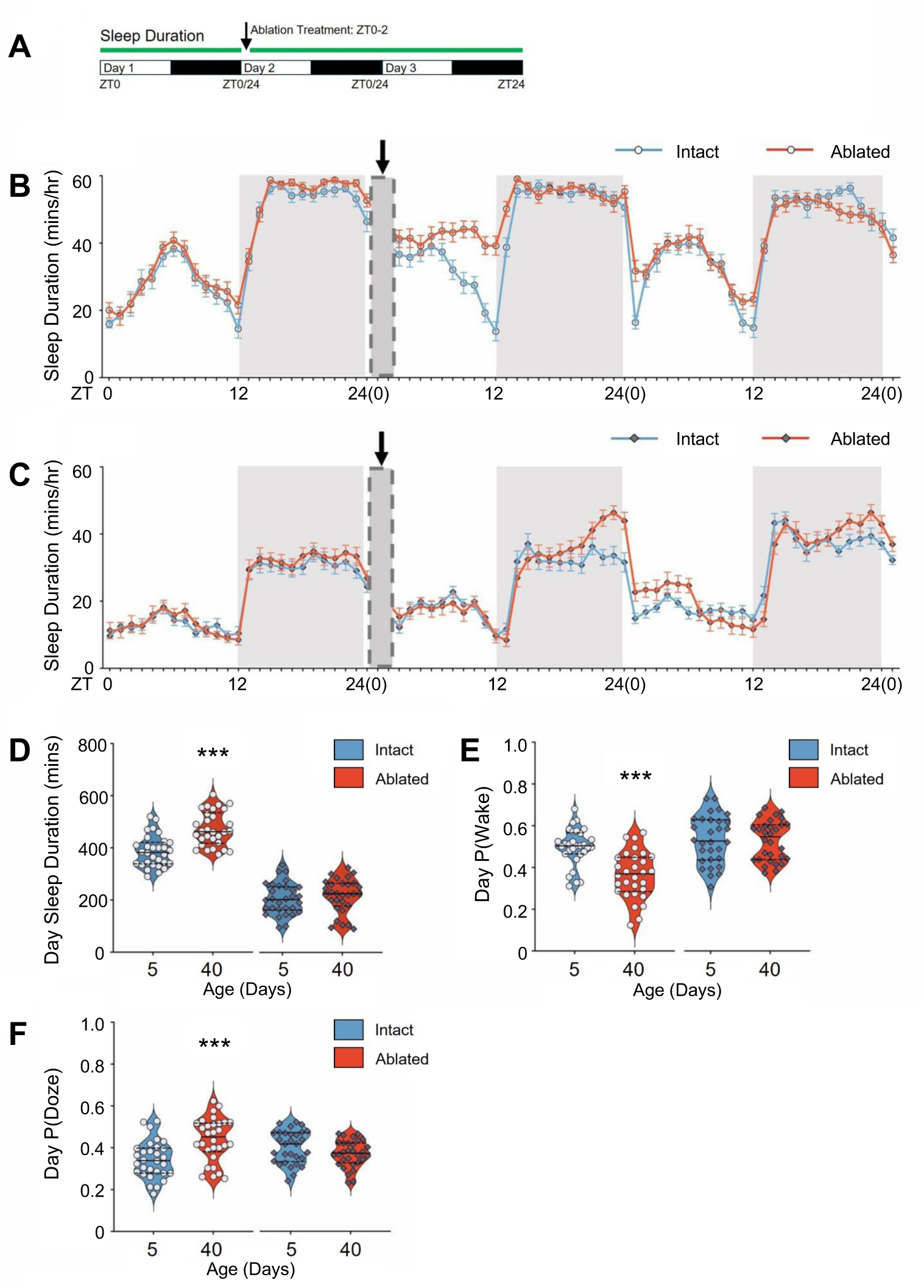
Antennal ablation-induced sleep response is absent in aged flies. (**A**) Diagram of experimental design for sleep response to neuronal injury in young (5-day-old) and aged (40-day-old) flies. (**B**,**C**) Sleep profiles prior to and following axonal injury in control unablated and ablated flies at (**B**) 5 days and (**C**) 40 days of age. Arrow denotes the time of antennal ablation, with the grey box (ZT0-2) representing a subsequent one-hour recovery period. White and gray boxes denote daytime and nighttime. (**D**) Daytime sleep duration was significantly increased in ablated flies following injury at 5 days of age but not at 40 days (P=0.0002 and P=0.4956, respectively). (**E**) Daytime wake probability, P(Wake) was significantly reduced in 5-day-old flies but not in 40-day-old flies (P<0.001 and P=0.8644, respectively). (**F**) Daytime sleep probability, (P)Doze, was significantly increased in 5-day-old flies following axotomy but not in 40-day-old flies (P=0.0003 and P=0.0843, respectively). n=30-35. *P<0.05; **P<0.01; ***P<0.001. Error bars indicate ± SEM.

We applied a Markov model that determines propensity to remain asleep (pDoze), an indicator of sleep depth [40]. Sleep propensity (pDoze) was elevated and wake propensity (pWake) was reduced in 5-day-old flies, but not 40-day-old flies, following injury, supporting the notion that sleep drive is lost in aged flies (Fig 2E,F; Fig S2B-C). Further, ablation induced an increase in average bout length during the day but not night in 5-day-old, but not 40-day-old flies (Fig S2F-G, S2J-K). Therefore, aging disrupts sleep-induction following injury. These findings support the notion that both reduced levels of basal sleep, and impaired sleep induction following injury contribute to age-related loss of glial activation and Wallerian degeneration.

The engulfment phenotypes observed in aged flies phenocopy those previously reported in response to acute sleep deprivation, raising the possibility that sleep deficits contribute to the age-related decline in glial engulfment [30]. To examine whether age-associated sleep loss contributes to loss of ensheathing glial response to injury, we pharmacologically enhanced sleep in young and old flies following antennal ablation and measured the effects on glial plasticity and axon engulfment [36,41]. Five- or 40-day old flies were fed the sleep-inducing GABA agonist gaboxadol for 48 hours following injury, followed by the quantification of Wallerian degeneration (Fig 3A; Fig S3A) [41]. Gaboxadol treatment significantly increased sleep in both young and old flies by increasing the average bout length (Fig 3B,C; Fig S3B,C). Signal from Or22a::GFP-labeled neurites was significantly reduced in ablated 5- and 40-day-old flies treated with Gaboxadol compared to solvent treated controls revealing that pharmacological induction of sleep restores glial engulfment to old flies (Fig 3D,E; Fig S3D,E). In both 5- and 40-day-old flies there was no effect of gaboxadol treatment on unablated controls, confirming that gaboxadol does not promote axonal engulfment in uninjured flies controls (Fig 3F; Fig S3F). Similarly, Draper levels were elevated in both 5 and 40-day-old ablated flies treated with gaboxadol compared to age-matched untreated controls, while there was no effect of gaboxadol on Draper levels in flies with intact antennae (Fig 3F’,G’,I; Fig S3F’,G’,I). Together, these findings reveal that sleep restores age-related deficits in neurite clearance and glial activation.

**Figure 3.**
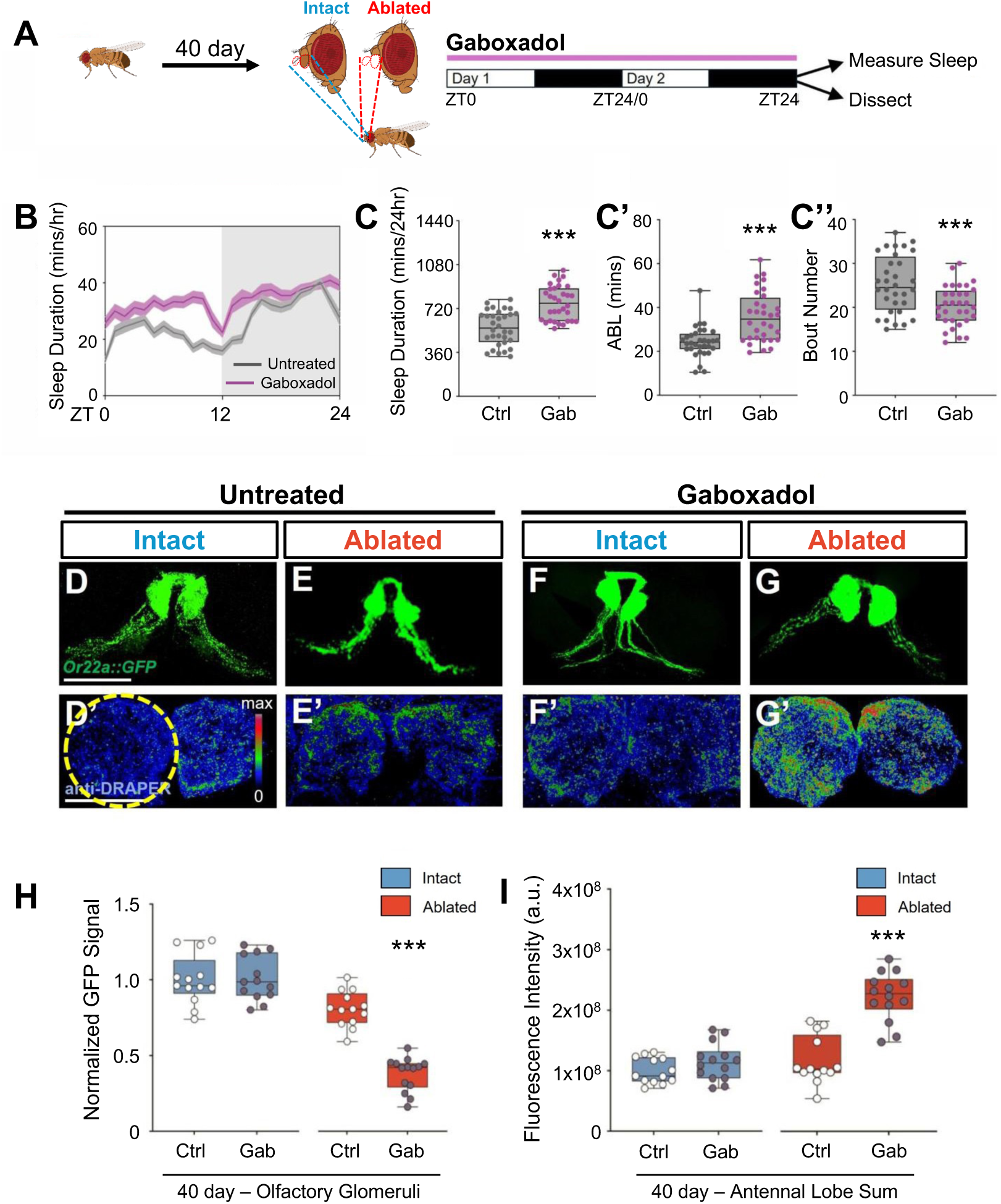
Gaboxadol treatment restores glial engulfment in aged flies. (**A**) Diagram of gaboxadol treatment and subsequent behavior monitoring and sample preparation. (**B**) Sleep profile of 40-day control and gaboxadol treated flies. (**C**) Total sleep and (**C’**) average bout length was significantly increased by gaboxadol treatment in 40-day-old flies (P<0.0001 and P<0.0001, respectively). while (**C’’**) bout number significantly decreased (P<0.001). n=30-35. (**C**) Total sleep and (**C’**) average bout length was significantly increased by gaboxadol treatment in 40-day-old flies (P<0.0001 and P<0.0001, respectively). while (**C’’**) bout number significantly decreased (P<0.001). (**D-G**) GFP signal labeled by Or22a::GFP and (**D’-G’**) immunostaining for DRAPER in the antennal lobes in gaboxadol treated and untreated intact or axotomized 40-day-old flies. Scale bar denotes 50 μm. (**H**) GFP quantification revealed no difference between gaboxadol treated (dark dots) and control (white dots) intact flies (P=0.8833). There was a significant reduction in GFP intensity in ablated gaboxadol treated flies compared to all other groups (P<0.001). (**I**) Draper levels were significantly elevated in ablated 40-day-old gaboxadol treated flies compared to all other groups (P<0.001). n=13-15. ***P<0.001 by one-way ANOVA followed by Turkey’s post-hoc tests. Error bars indicate ± SEM.

It is possible that the effects of gaboxadol on glial activation and neurite engulfment are due to activation of GABA signaling rather than induction of sleep per se. To differentiate between these possibilities, we genetically activated R23E10 neurons that label sleep promoting neurons in the brain and/or the ventral nerve cord [42–44] and measured the effect on engulfment of damaged neurite s. Briefly, we ablated the antennae of flies expressing the thermosensitive cation channel *TrpA1* in R23E10 neurons (R23E10-GAL4>TrpA1), and increased the temperature from 22°C to 29°C for 48 hours following injury. Following sleep induction, brains were immunostained for Draper levels (Fig 4A). Consistent with previous findings in young flies, thermogenetic activation of 23E10 neurons increased sleep in 5- and 40-day-old flies (Fig 4B-D; Fig S4A-E) [42]. Induction of sleep enhanced Draper levels in 5-day-old flies compared to genetic controls (Fig 4E,F). There was no effect of sleep induction on Draper levels in intact 5-day-old flies, suggesting the effects of sleep on Draper is specific to post-injury response (Fig 4F). Similarly, in 40-day-old flies, genetic induction of sleep increased Draper levels in ablated flies but not in unablated controls (Fig 4G,H). Together, these findings confirm that genetic induction of sleep is sufficient to restore post-injury Draper induction in old flies.

**Figure 4.**
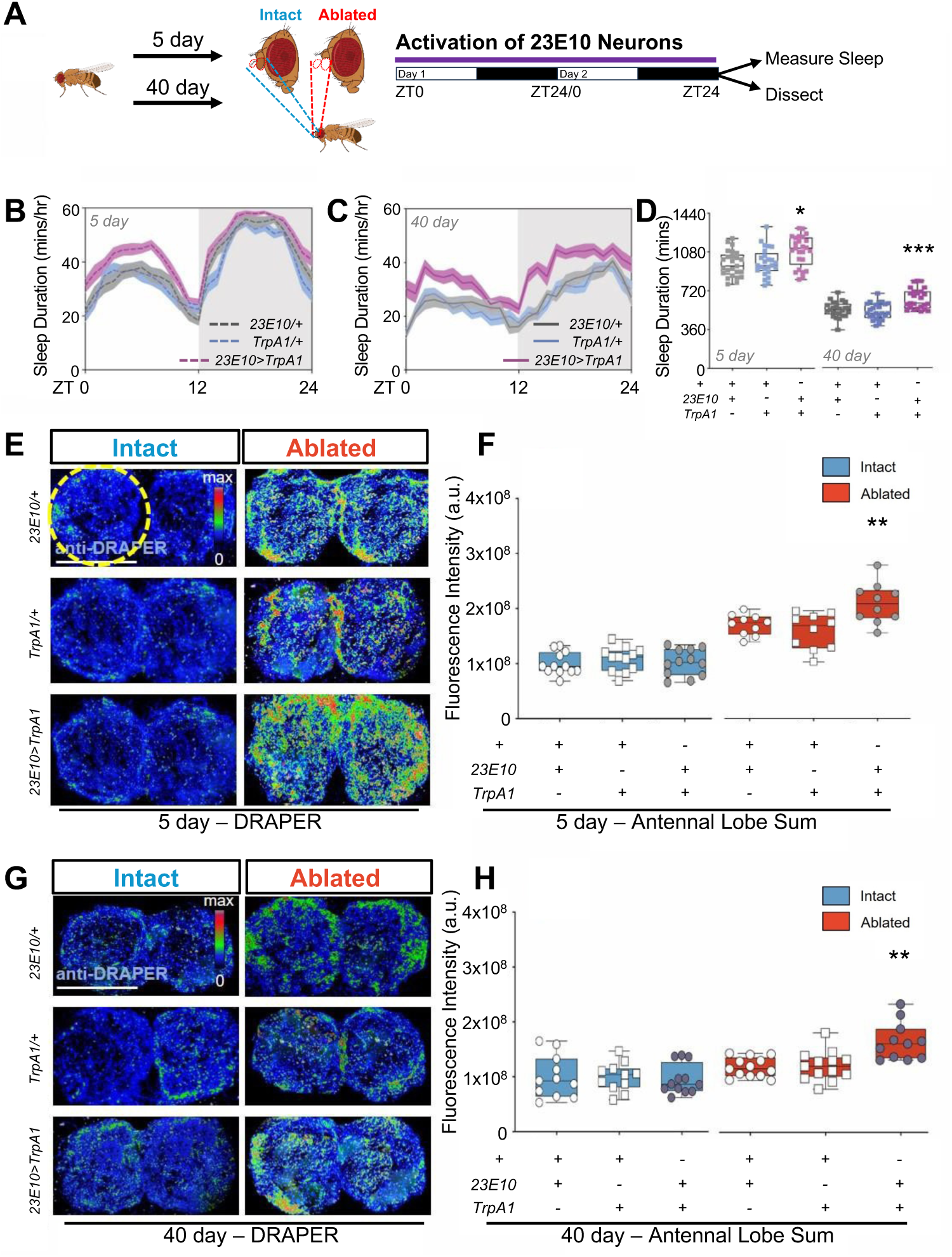
Robust Draper recruitment restores glial engulfment in aged flies through genetic induction of sleep. (**A**) Diagram of sleep induction by thermogenetic activation of 23E10 neurons and subsequent sleep measurement and quantification of glial engulfment. (**B**-**D**) Elevated temperature significantly induced sleep in 5-day-old (**B**) and 40-day-old (**C**) 23E10-GAL4>TrpA1 flies compared to TrpA1/+ or 23E10-GAL4/+ controls. (**D**) Total sleep was significantly increased after two-day thermo-activation of 23E10 neurons in both 5-day-old (P=0.0022) and 40-day-old (P=0.0001). n=30-35. (**E**) Immunostaining for Draper levels in the antennal lobes in control (23E10/+,TrpA1/+) and experimental (23E10>TrpA1) 5-day-old flies. (**F**) There were no differences in Draper levels in intact flies (P=0.8455), while Draper levels were significantly elevated in 23E10>TrpA1 ablated flies, compared to ablated flies without sleep induction (P=0.0072). (**G**) Immunostaining for Draper levels in the antennal lobes in control (23E10/+,TrpA1/+) and experimental (23E10>TrpA1) 40-day-old flies. (**H**) There were no differences in Draper levels in intact flies (P=0.7409), while Draper levels were significantly elevated in 23E10>TrpA1 ablated flies, compared to ablated flies without sleep induction (P=0.0003). n=10-14. Scale bar denotes 50 μm. *P<0.05, **P<0.01, ***P<0.001. Error bars indicate ± SEM.

Numerous genetic pathways have been identified as contributing to glial activation and increased Draper levels following axotomy, yet much less is known about how aging impacts glial function [8]. To identify age-dependent changes in ensheathing glia we performed single-nucleus RNA-sequencing (snRNA-seq) on the central brains of both young and old flies under intact and axotomized conditions (Figure 5A). We dissected 120-140 central brains of intact or ablated flies at 5 and 40 days of age. This yielded at least 295k nuclei for each of the four experimental groups, resulting in 68,361 nuclei that met the criteria for inclusion in analysis (see methods). There were no significant differences in nuclei number between any of the groups tested, suggesting consistency between sample collections. We were able to classify nuclei into diverse types of brain cell types based on known markers (Fig S5A-D). Clustering analysis revealed six distinct populations including cortex glia, ensheathing glia, astrocyte-like glia, perineural glia, subperineural glia and chiasm giant glia, and one unannotated group. (Fig 5B-D and Fig S5E). The total number, and types of glia did not vary significantly across experimental conditions, supporting the notion that loss of neurite engulfment in aged flies is not due to an overall loss of ensheathing glia (Fig 5E, S5F).

**Figure 5.**
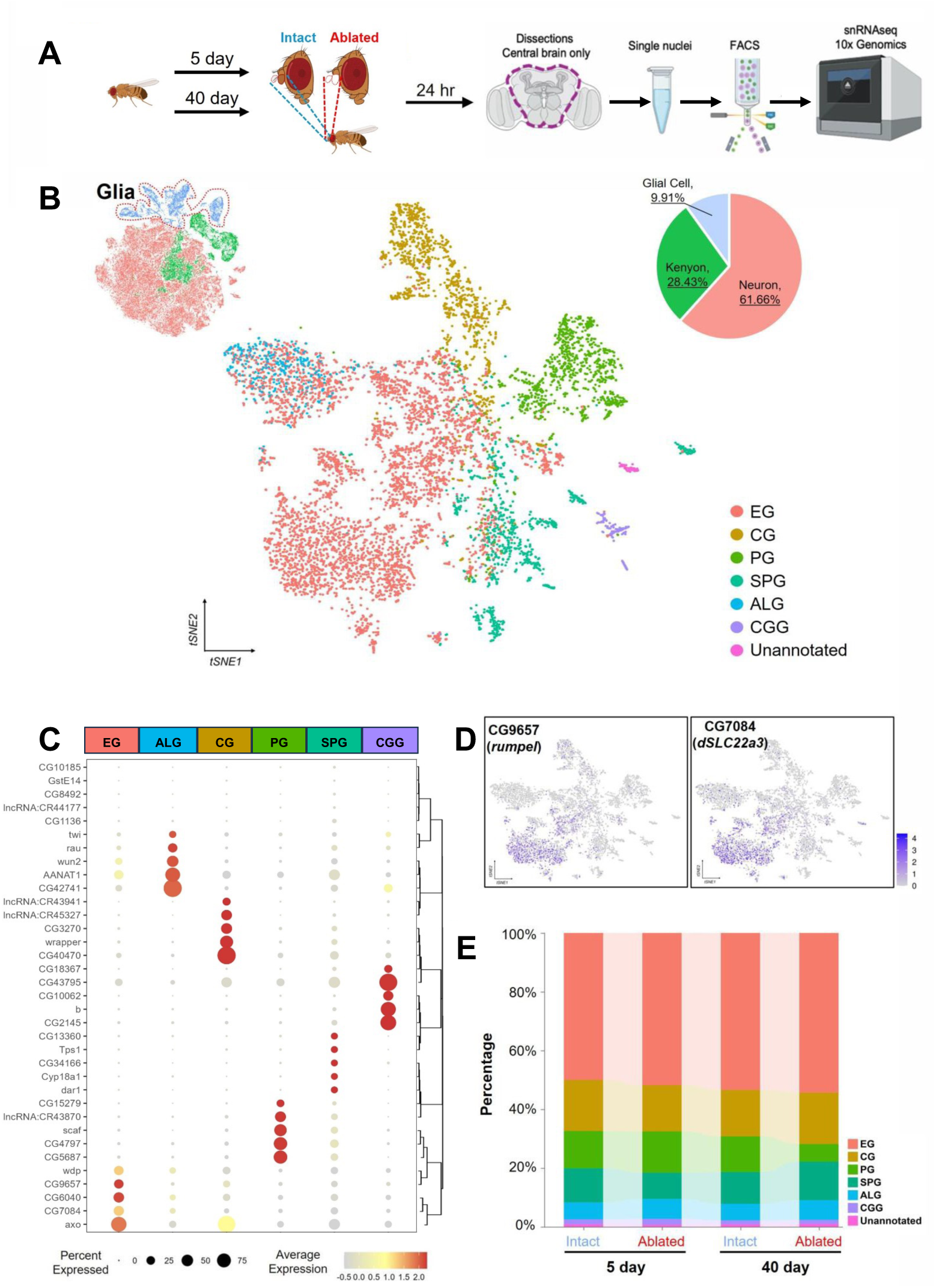
snRNA-sequencing of central brains in young and aged intact or ablated flies. (**A**) Schematic of tissue collection and snRNA-sequencing workflow. Axotomy of the third segment of antenna was performed for the adult flies at 5- or 40-days of age. After 24 hr of recovery following the axotomy, the central brains of intact or ablated flies were dissected and dissociated to collect single nuclei. (**B**) t-distributed Stochastic Neighbor Embedding (t-SNE) visualization of the 7 distinct subclusters of glial cells from the integrated nuclei of 5-day intact/ablated and 40-day intact/ablated pooled replicate samples. Each color corresponds to a unique cluster, and each dot represents a single nucleus. EG, ensheathing glia; CG, cortex glia; PG, perineurial glia; SPG, subperineurial glia; ALG, astrocyte-like glia; CGG, chiasm giant glia; Unannotated, unannotated glial types. (**C**) Dotplot of marker genes for each glial subtype that were used for annotation. (**D**) t-SNE visualization of selective EG marker genes, *rumpel* (left) and *dSLC22a3* (right), on glia cohort. (**E**) Percentage of glial subtypes among four group samples.

To identify genes that are transcriptionally regulated in response to neural injury in young and old flies, we compared gene expression across experimental groups within the ensheathing glia cluster. We first sought to specifically identify the olfactory ensheathing glia based on the expression of *repo* and the Toll-1 receptor *dSARM*, and *Draper*, all of which are highly expressed in olfactory ensheathing glia post neural injury[45–47]. We specifically analyzed the ensheathing glia group. Of these, 16.31% co-expressed *Draper* and *Sarm*, and were therefore defined as olfactory ensheathing glia (Fig 6A and S6A). This subset was further examined for differential gene expression.

**Figure 6.**
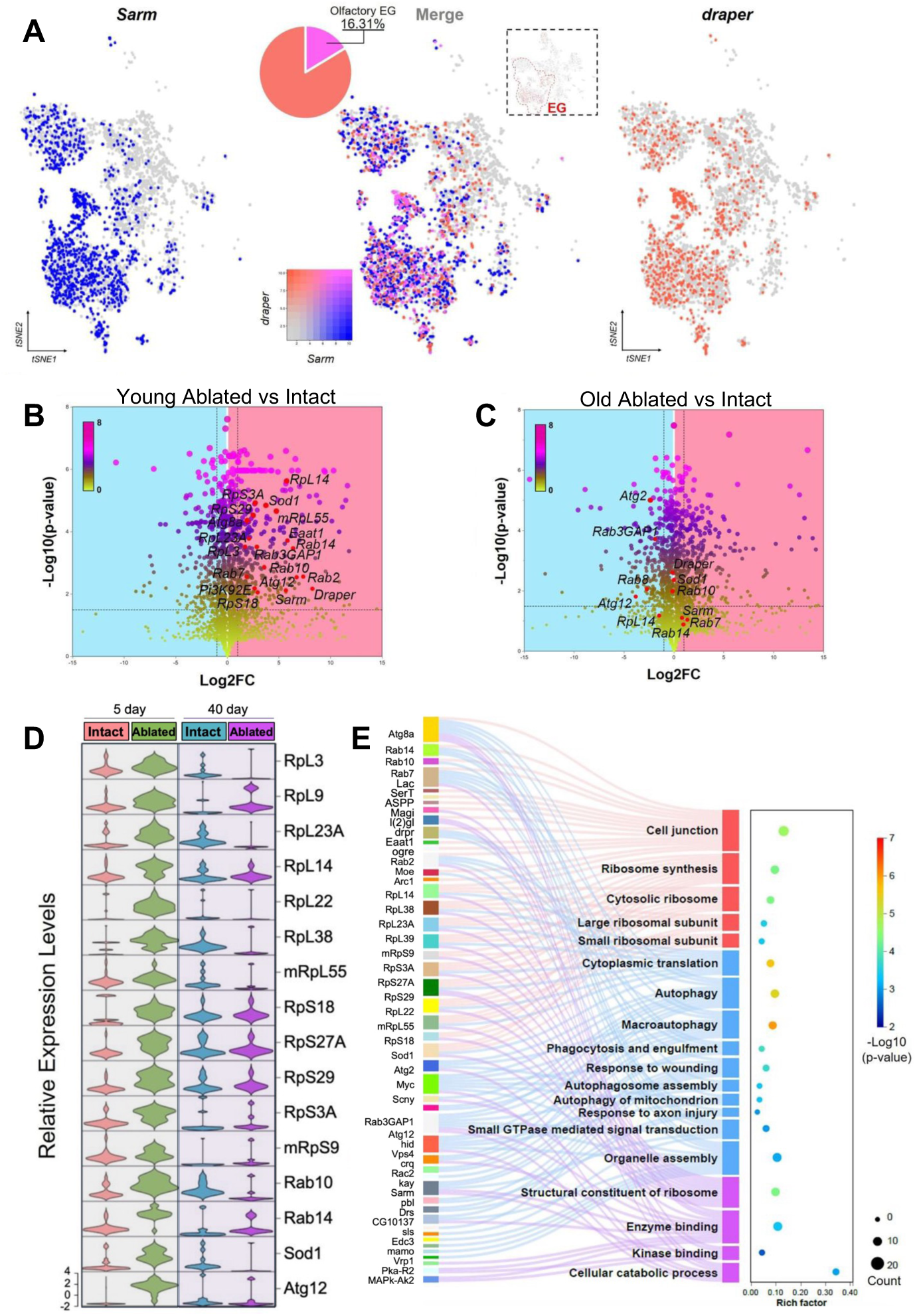
Differential gene expression analysis of *Sarm* and *draper* positive olfactory ensheathing glia in young and aged intact or ablated flies. (**A**) t-SNE visualization of *Sarm* and *draper* positive olfactory ensheathing glial nuclei. Nuclei that express *Sarm* (*Sarm*^+^) are highlighted in blue, while nuclei that express *draper* (*draper*^+^) are highlighted in red. *Sarm* and *draper* co-expressed olfactory EG were merged in pink with a color box indicated the expression levels of *Sarm* or *draper* in ensheathing glial cell. **(B)** Volcano plot showing the differential gene expression in *Sarm^+^* and *draper^+^* olfactory ensheathing glial nuclei between 5-day ablated and intact treatments. (**C**) Volcano plot showing the differential gene expression in *Sarm^+^* and *draper^+^* olfactory ensheathing glial nuclei between 40-day ablated and intact groups. The dashed lines represent the thresholds for significance. Genes with significant differential expression were highlighted in dark colors, while non-significant genes are shown in lime green with blue background for downregulated genes and pink background for upregulated (activated response to neural injury) genes. Promising candidate genes were highlighted in red dots. (**D**) Gene expression levels of candidate genes across each treatment. (**E**) Ribosomal protein subunit genes and autophagy and phagocytosis replated pathways were significantly enriched in 5-day-old flies post neural injury.

To determine the differences in gene expression between young and old individuals in response to neuronal injury, we compared the transcriptional changes in genes between the young and old groups of olfactory ensheathing glial nuclei, using Seurat’s FindMarkers function. In the comprehensive gene expression analysis, we observed distinct patterns of differential gene regulation between intact and ablated groups in young and old cohorts separately. In young individuals, the ablated group exhibited a notable divergence, with 927 genes significantly upregulated and 670 genes significantly downregulated compared to the intact group (Fig S6B). This robust response suggests a pronounced compensatory mechanism for engulfment of damaged neurites or altered developmental trajectory in response to ablation. Conversely, in aged individuals, the ablated flies displayed a distinct profile, with 877 genes significantly upregulated, of which 325 overlapped with those upregulated in the young group. This overlap indicates a subset of genes that maintain their function throughout the lifespan in response to neuronal injury (Fig S6B). However, among the 602 uniquely upregulated genes in the young ablated group, there were some key candidates such as autophagy related genes *Atg 12*, *Atg 8a*, *Rab 10*, *Rab 14*, *Rab 7* and *Rab3GAP1*, and a set of ribosome subunit genes, as well as *Draper* and *Sarm*, which may play pivotal roles in the efficient neurite engulfment within 24 hours post injury and warrant further investigation for their functional implications in development and aging (Fig 6B-C).

Next, we performed pseudobulk differential expression analysis on olfactory EG cell cluster from the raw count data and subsequently used z-score clustering expression to identify different patterns of gene expression across all four experimental groups (Fig S6C). We identified 1356 genes in cluster 2 that were selectively upregulated in young flies following ablation but not old flies. Through these two different analyses and overlapping the 602 gene list with genes in cluster 2, we specifically focused analysis on the genes that were upregulated in the young cohort, but not in aged flies post neural injury (Fig S6B-D). These candidate genes included a set of ribosome subunit genes, and some autophagy genes related to phagocytosis and engulfment pathways, all fall into this expression pattern (Fig 6D-E, S6D). Strikingly, 7 large ribosomal protein subunit (RpL) and 5 small ribosomal subunit protein (RpS) transcripts were significantly upregulated following ablation in young flies, but not old flies (Fig 6D-E). These genes provide numerous potential candidate regulators of glial engulfment and injury response and highlight a role for an increase in ribosomal subunits following injury. Gene ontology analysis revealed an enrichment for multiple aspects of ribosome function, as well as autophagy, suggesting this group known and novel cellular processes associated with Wallerian degeneration. Notably, we cannot deny that other glial subpopulations, apart from olfactory ensheathing glia, are not involved in this process, as some cells from other glial subtypes also co-expressed *Sarm* and *draper* after being ablated (Fig S6E-H).Together, these analyses identify numerous genes that are transcriptionally regulated by both neural injury and aging, providing candidate regulators for age-dependent loss of Wallerian degeneration.

## Discussion

Here, we show a link between reduced sleep and age-dependent loss of Wallerian degeneration in *Drosophila*. Both aging and acute sleep loss are associated with reduced brain function including changes in structural plasticity, long-term potentiation, memory, and elevated oxidative stress levels [25,27,31,48]. Previous work in mice and flies has shown that both aging and sleep loss impair glial engulfment, but the relationship between sleep loss and aging has been unclear[29,30]. Here, we show that pharmacological and genetic induction of sleep restores induction of Draper and glial-mediated engulfment to aged flies. These findings reveal a critical role for sleep-loss in age-dependent loss of glial function.

Age-related loss of glial function is associated with many aspects of declining brain function. In flies and mammals, the clearance of debris and toxic substances from the brain is critical for healthy aging. In mammals, both the overactivation and reduced function of microglia is associated with neurodegeneration[24,49,50], while the glymphatic systems play a critical role in regulating the clearance of amyloid beta from cerebral spinal fluid [22]. In *Drosophila,* loss of the ensheathing glia is associated with lipid accumulation and reduced lifespan, while genetically increasing ensheathing glia number is protective in a model of Alzheimer’s disease [51]. Further, loss of glial function accelerates age-related changes in memory [52]. These findings suggest ensheathing glia are critical for healthy aging. Here, we find that age-related sleep loss contributes to reduced glial function. These findings raise the possibility that loss of glia function may contribute to the negative impacts of aging.

There are many functional parallels between *Drosophila* and mammalian glia and their roles in sleep regulation. Both astrocytes and microglia are essential for normal brain function in animals ranging from flies to mice. These effects appear to be highly conserved because astrocytic-like and microglia-like ensheathing glia also regulate sleep in *Drosophila* [16,53,54]. In some cases, glia appear to impact global brain function, such as regulation of synaptic strength and neural excitability, while in other cases glia appear to modulate levels of specific sleep transmitters or neural circuits [55–58]. It is possible that the effects of sleep and aging in modulating glia function extend beyond olfactory ensheathing glia, to other types of glia in the brain. Future studies examining the effects of aging on astrocyte-like, cortex glia, and non-olfactory ensheathing glia is critical to understand whether aging and sleep loss impact all types of glia or are specific to the engulfing properties of olfactory ensheathing glia.

*Drosophila* increase sleep immediately following ablation injury to the antennae or wing suggesting this is a naturally occurring mechanism to promote recovery and glial-mediated engulfment[18]. The degree of post-injury sleep is thought to directly result from synaptic pruning, because expression of neuroprotective factors in the olfactory neurons blunt sleep following injury [18]. Here, we show that post-injury sleep is reduced in aged flies, consistent with the notion that pruning promotes sleep following injury. These findings raise the possibility that bi-directional interactions between neurite clearance and sleep are both impacted by aging. The effects were greatest in the first 12 hours following injury. These findings raise the possibility that greater upregulation of neuroprotective factors contribute to the blunted sleep response. Because ORN cell bodies are removed in the process, we are unable to address this using our sn-RNAseq protocol. However, future work specifically examining interactions between post-injury sleep induction and neuroprotective factors in aged flies may be informative.

Transcriptional regulation in glia is modulated by sleep, circadian timing, and age [59,60]. These molecular factors directly impact glial function including engulfment of debris, regulation of neurotransmitter release, and secretion of sleep-regulating factors [54,61]. For example, insulin signaling promotes sleep and activates ensheathing glia phagocytotic mechanisms in Drosophila, while diet induced insulin resistance inhibits neurite clearance, revealing a critical role for insulin signaling in debris clearance [13,28]. To identify new factors that contribute to age related changes in sleep we performed sn-RNAseq in the brains of old and young flies with ablated antennae. We find numerous genes previously known to be under transcriptional regulation including *Draper* and the autophagy regulators *Atg8a* and *Rab5* are not upregulated following axotomy in aged flies. Altered autophagy is associated with sleep, with autophagosome levels accumulating during periods of wakefulness, and sleep loss is associated with reduced proteostasis and ultimately neurodegeneration [62,63].

Our findings that aging inhibits induction of ribosomal subunit transcription in glia is consistent with studies ranging from *C. elegans* to humans revealing age-associated loss of ribosomal protein subunits [64,65]. Therefore, the loss of ribosomal biogenesis may inhibit the structural and functional plasticity required to engulf aging neurites. It is possible that the injury-induced ribosomal subunits are generally required for enhanced translation following injury, or that the subunits confer unique functions. In *Drosophila*, selective loss of individual ribosomal subunits is associated with defects in development and cellular function [66–68] Furthermore, the TOR pathway has been implicated more generally in age-associated loss of ribosomal biogenesis including reduced TOR signaling [69]. Therefore, it is possible that pharmacological enhancement of ribosomal biogenesis through these pathways has potential to ameliorate the cellular, and possibly functional, effects of age-related sleep loss [70].

Taken together, our findings reveal a role for sleep in age-related loss of glial-mediated phagocytosis. Sleep is sufficient to restore glial membrane plasticity, induction of Draper, and neurite engulfment in aged flies. Our findings provide a system to examine the effects of sleep on glial-mediated phagocytosis and broader aspects of how sleep and aging impact glial function. Given the many aspects of glial function, including roles in regulating many aspects of brain health and function, are conserved from flies to mammals, this system has potential to advance our understanding of changes in glial function across the lifespan.

## Materials and Methods

### Fly stocks and maintenance

Flies were reared and maintained on a 12:12 light–dark cycle in humidified incubators at 25°C and 65% humidity (Percival Scientific, Perry, IA, USA). The following fly lines were obtained from the Bloomington Stock Center: UAS-mCD8GFP (#32186; [71]) *Repo*-GAL4 (#7415; [72]), OR22a::GFP (#52620; [73]), *23E10*-GAL4 (#49032); UAS-*TrpA1* (#26263; Paul Garrity). The background control line used in this study was *w^1118^* (BL# 5905; [74]) and all experimental stocks were either generated in this background or outcrossed to *w^1118^* for a minimum of six generations prior to testing. Unless indicated otherwise, mated females reared on standard *Drosophila* food media (Bloomington Formulation, Nutri-fly, #66-113, Genesee Scientific) were used for all experiments. For aging experiments, flies were isolated 1 day post-eclosion, sorted by sex into vials of ∼30 flies, and then transferred to new vials three times per week until they reached the indicated age. Unless otherwised noted, all experiments were performed on male flies.

### Fly axotomy

Fly axotomoy was performed as previously described[30]. Flies of the indicated age and genotype were gently anesthetized on CO_2_ for less than one minute at ZT0-2. Next, using forceps, the third segments of both antennae were removed. Controls received the same treatment without ablation. Unless indicated otherwise, control and ablated flies were maintained on standard food for 24 hours prior to dissection.

### Drug treatment

Gaboxadol (4,5,6,7-retrahydroisoxazolo[5,4-*c*] pyridin-3-ol hydrochloride, THIP hydrochloride; Sigma Aldrich, St. Louis, MO, USA, #T101) was dissolved in dH_2_O at 1 mg/ml, then mixed to 0.01 mg/ml in cooled, standard fly food, as previously described [41,75]. Axotomized and intact control flies were placed on gaboxadol-laced food immediately following axotomy, where they remained for the duration of the experiment.

### Measurements of sleep behavior

Sleep behavior was measured using the *Drosophila* Activity Monitoring (DAM) system (Trikinetics, Waltham, MA), which detects activity by monitoring infrared beam crossings of individually housed flies. The number of beam-breaks is used to determine the amount of sleep, along with associated sleep metrics, by identifying 5-min bouts of quiescence using the *Drosophila* Sleep Counting Macro [76,77]. All experiments were performed in incubators at the same temperature and humidity mentioned above. For sleep experiment in aging flies, flies of the indicated age were briefly anesthetized with CO_2_ and loaded individually in DAMs tubes with standard fly food. Flies were acclimated for a minimum of 24 hours prior to the start of behavioral analysis. Measurements of sleep were then measured over a 24 hr period starting at ZT0. For sleep experiments in axotomized flies, sleep was measured for 24 hr as described in aging flies, however at ZT 0-2 the following day, individual flies were removed, axotomized, and then returned to their respective tube and monitor, in which sleep was recorded for the subsequent 48 hours post axotomy. Control flies received the same treatment without ablation. For sleep-induction experiments, sleep was measured in flies fed 0.01 mg/ml Gaboxadol. For 23E10-GAL4 activation experiments, sleep was measured at the permissive temperature of 29°C.

### Immunohistochemistry and imaging

Brains of the indicated age and treatment were dissected in cold phosphate buffer solution (PBS) and fixed in 4% paraformaldehyde in PBS containing 0.5% Triton-X100 [PBT], for 40 minutes at room temperature (RT). After three washes (15 min each) with PBT and rinsing overnight at 4°C, the brains were incubated with relevant primary antibody, anti-GFP (1:1500; #PA1-980A, Fisher Scientific) or anti-DRAPER 5D14 (1:500; #draper, AB_2618105, Developmental Studies Hybridoma Bank), for 48 hr at 4°C with rotation. After three 15-min washes in PBT at RT, the brains were incubated with secondary antibody for 90 min at RT: (Alexa Fluor 555 donkey anti-mouse IgG, Invitrogen, Carlsbad, CA, USA, #A31570; Alexa Fluor 488 goat anti-rabbit IgG, Invitrogen, Carlsbad, CA, USA, #A11008), and then washed three times (30 min each) in PBT, followed by a final overnight wash in PBT. Brains were mounted in Vectashield (H1000, Vector Labs, Burlingame, CA, USA). Fluorescence images were acquired on a Nikon A1R laser-scanning confocal microscope. Brains were imaged using the 561 nm and 488 nm lasers sequentially, with imaging settings (e.g., laser power, gain, and zoom) kept constant throughout the entire experiment. Image stacks were acquired using the ND Sequence Acquisition function, imaging from the above to below the antennal lobes using a 2-μm step size.

### Quantification of fluorescence intensity

Quantification of *OR22a::GFP* fluorescence intensity was performed by generating a sum intensity projection of the *OR22a*-labeled neurons and quantifying the sum fluorescence GFP intensity using the Nikon AR Image Analysis software (Nikon, Melville, NY, USA), as previously described [9]. *Repo-*GAL4*>*UAS-mcD8GFP, *MZ0709*-GAL4>UAS-mCD8GFP and anti-DRAPER were quantified using a standardized ROI of the outer, dorsal-medial antennal lobe membrane. The mean of the intensity averaged from each slice was used. Images are presented as the Z-stack projection through the entire brain and processed using Fiji [78].

### Single nuclei sequencing

#### Tissue collection, nuclei isolation, library preparation, and sequencing

Central brains from either axotomized or intact flies were dissected at the age of 5 and 40 days and then placed into 1.5 mL RNase-free Eppendorf tubes, flash-frozen in liquid nitrogen, and then stored at -80°C. All dissections were performed between ZT0 and ZT4. Approximately 120-140 central brains were collected for each treatment and stored on dry ice. Nuclei isolation and snRNAseq were performed by Singulomics Corporation (Singulomics.com; New York). Briefly, tissue was homogenized and lysed with Triton X-100 in RNase-free water to isolate nuclei. The nuclei were then purified, centrifuged, resuspended in PBS with RNase Inhibitor, and then diluted to ∼700 nuclei/µL. Next, standardized 10x capture and library preparation were performed using the 10x Genomics Chromium Next GEM 3’ Single Cell Reagent kit v3.1 (10x Genomics; Pleasanton, CA). The prepared libraries were then sequenced using Illumina NovaSeq 6000 (Illumina; San Diego, CA). The snRNA-seq raw sequencing files were processed using CellRanger 6.0 (10x Genomics; Pleasanton, CA) and then mapped to the *Drosophila* reference genome (Flybase r6.31).

#### Data normalization and integration

All analysis of snRNA-seq data was performed in R (v4.3) using the package Seurat (v5.0.2; [79]). Unless otherwise noted, default parameters were used for all operations. Each sample was independently normalized using the SCTransform function (v2;) [80,81]. After normalization, integration features were selected using the SelectIntegrationFeatures function, with nFeatures = 3000. Normalized data was prepared for integration with the PrepSCTIntegration function. Integration anchors were identified using the FindIntegrationAnchors function, with normalization.method = “SCT”. Lastly, using these anchors, the data was integrated with the IntegrateData command.

Following integration, Principal Component Analysis was performed using the RunPCA function, with npcs = 30. The resulting PCs were used to identify clusters, with a resolution of 0.2, resulting in 27 total clusters. The integrated data was prepared for marker identification by running the PrepSCTFindMarkers function, and then markers were identified with the FindAllMarkers command, using the MAST statistical test [82], which has been found to perform well on single-cell data sets [83].

The list of marker genes was used to annotate cluster identity. First, we compared the marker genes for each cluster to previously generated datasets in *Drosophila*, using the Cell Marker Enrichment tool (DRscDB; https://www.flyrnai.org/tools/single_cell/web/enrichment [84]). Wherever possible, we refined our labels by comparing marker genes with published literature marking known cell types [85].

#### Differential gene expression analysis

To identify genes within nuclei that are differentially expressed between each of the different treatments, we applied the FindMarkers function on these cells, again using the MAST statistical test, and set group.by= ‘treatment’, separately in the you or old cohort. Genes with an adjusted *P* value less than 0.05 and an average Log_2_ Fold Change greater than 0.25 or less than -0.25 were considered differentially expressed and used for downstream analysis. To comprehensively compare gene expression within the same cell type across young and old samples under both intact and ablation conditions, as well as to validate the differential expression genes (DEGs) identified by the FindMarkers function, pseudobulk differential expression analysis was performed as needed. First we use AggregateExpression() function to sum together gene counts of all the cells from the same sample for each cell type. This involves generating matching metadata at the sample level for each cell type. Then we normalized the pseudobulk count matrix and performed DE analysis using DESeq2 package (v1.46.0) to identify genes that are differentially expressed between experimental conditions. Variance-stabilizing transformation (VST) was applied to the count data to stabilize the variance across different expression levels and makes the data more suitable for downstream analysis. To visualize gene expressions in different patterns, we normalized the expression data by calculating z-scores for each gene across samples and then used ClusterGVis package (v0.0.2) for clustering and visualizing gene expression trends.

#### Subclustering glia clusters

We first identified all clusters labeled as ‘Glia’ through UMAP analysis of our entire data set and extracted these cells for further analysis. Subsequently, PCA was conducted exclusively on these glial cells. The two principal components (PCs) that were found to be significant were then utilized to carry out a new round of dimensionality reduction, which was guided by Jackstraw analysis. The resolution for clustering was set at 2.5, which was determined by visually inspecting the distinct UMAP populations and ensuring that they could be clearly identified as separate clusters. Finally, differential expression analysis was performed between each of the different treatments in each glial subtype clusters, employing the same FindMarkers function as previously described.

#### Gene Ontology Analysis

To identify the overarching GO terms of the resulting lists of significant ablation markers, we used clusterProfiler package (v4.8.3), as custom background, to conduct gene set enrichment analyses. The log2 fold change (log2FC) values were calculated for each gene expressed in the cell population of interest, comparing the young ablated and intact groups. The sorted log2FC list was then input into the gseGO function with the following parameters: ontology set to ‘Biological Process’ (ont = ‘BP’), gene identifier type set to ‘SYMBOL’ (keyType = ‘SYMBOL’), minimum gene set size set to 3 (minGSSize = 3), maximum gene set size set to 800 (maxGSSize = 800), organism database set to org.Dm.eg.db (OrgDb = org.Dm.eg.db), and p-value cutoff set to 0.01 (pvalueCutoff = 0.01). GO terms were subsequently grouped according to parent terms based on semantic similarity as calculated using the mgoSim() function of the GOSemSim R package (v.2.20.0) and the reduceSimMatrix() function in the rrvgo package (v.1.6.0)[86].

### Statistical Analysis

All data of sleep duration and florescence intensity are presented as box plots with ± SEM error bar. Unless stated otherwise, A one-way analysis of variance (ANOVA) was employed for comparisons involving two or more genotypes subjected to a single treatment, whereas a two-way ANOVA was utilized for comparisons involving two or more genotypes subjected to two treatments. Post hoc analyses were conducted using Sidak’s multiple comparisons test. Each experiment was conducted with a minimum of two independent runs, and the sample size for each genotype and treatment combination is detailed in the respective figure legends. All statistical analyses were performed using InStat software GraphPad Prism (version 10.3.0 for Windows, Boston, Massachusetts USA), where N denotes the number of biological samples tested.

## Acknowledgements

The authors are grateful to members of the Keene Lab for technical assistance and helpful discussion. This manuscript was supported by the National Institutes of Health (NIH) grants R01NS131628 and R01HL143790 to ACK and <u>R00AG071833</u> to EBB. The authors thank Dr. Akhila Rajan (Fred Hutchinson Cancer Institute) and James Cai (Texas A&M) for helpful suggestions and feedback.

## Author Contributions

ACK designed experiments, with guidance from IF. J.Z conducted experiments; J.Z and E.B.B. and E.L. conducted data analysis. J.Z. designed figures. All authors played a role in designing experiments, interpreting data, and writing the paper.

## Declaration of interests

The authors declare no competing interests.

## Supplementary Figures

**Supplementary Figure 1.**
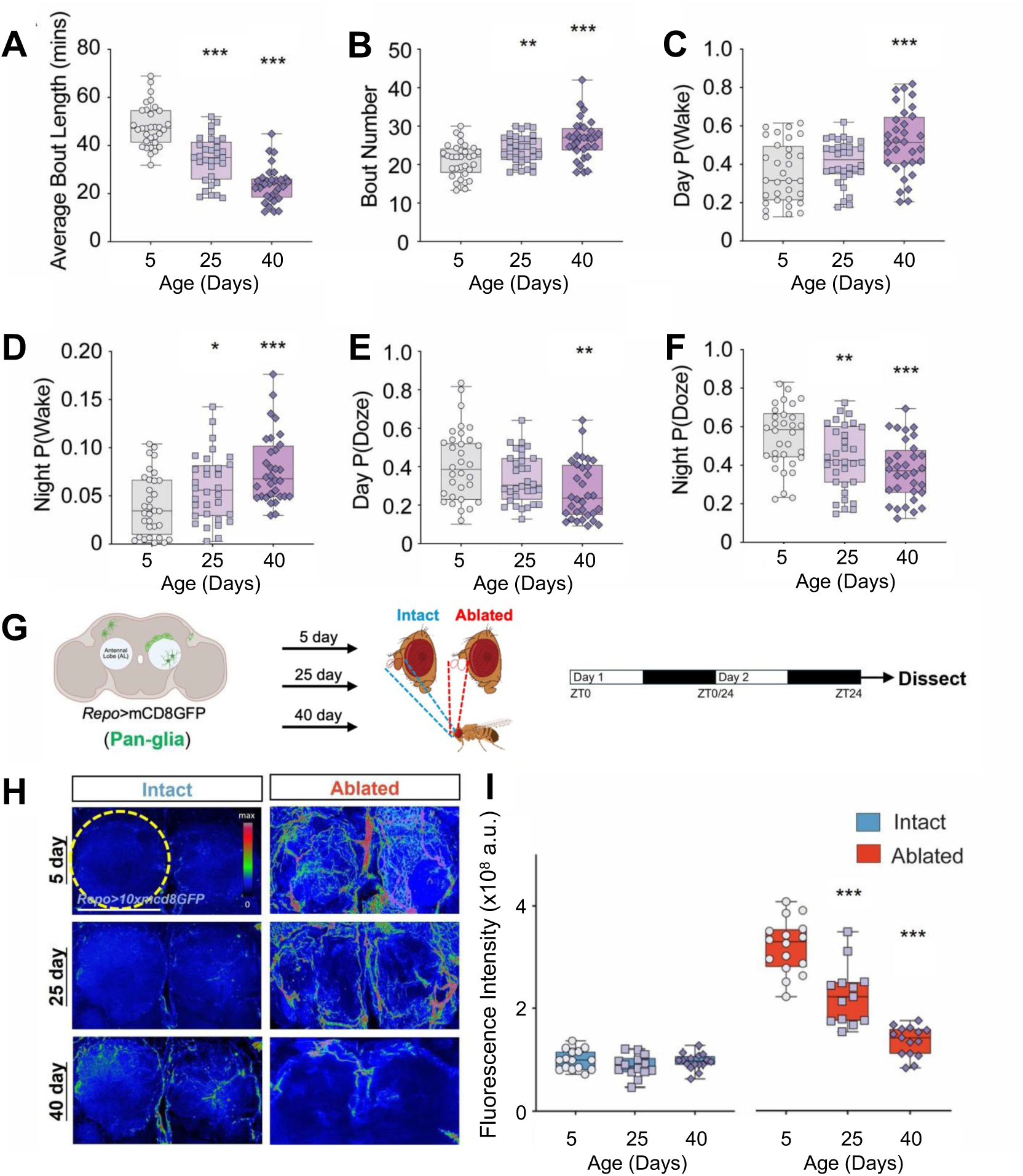
Pan-glial innervation response to neuronal injury is reduced in aged flies. (**A**,**B**) Reduced average bout length (ABL) and increased bout number in 25 day and 40 day indicated sleep became much more fragmented as flies aged. (**C,D**) Wake probability, P(Wake), significantly increased with age both during the (**C**) day and (**D**) night (P<0.001 and P<0.001, respectively). (**E,F**) Sleep probability, P(Dose), significantly decreased with age both the (**E**) day and (**F**) night (P=0.0087 and P<0.001, respectively). **(G**) Schematic of experimental design for pan-glia response to neuronal injury in young (5-day-old) and aged (25-day-old or 40-day-old) flies. (**H**) Pan-glial innervation of the antennal lobes in *Repo*-GAL4>mCD8GFP flies at 5-, 25- and 40-days of age. Scale bar denotes 50 μm. (**I**) Pan-glial response post injury is significantly reduced in ablated flies at 25-(P<0.001) and 40-days of age (P<0.001). Tukey’s multiple comparison tests: *P<0.05; **P<0.01; ***P<0.001. Error bars indicate ± SEM.

**Supplementary Figure 2.**
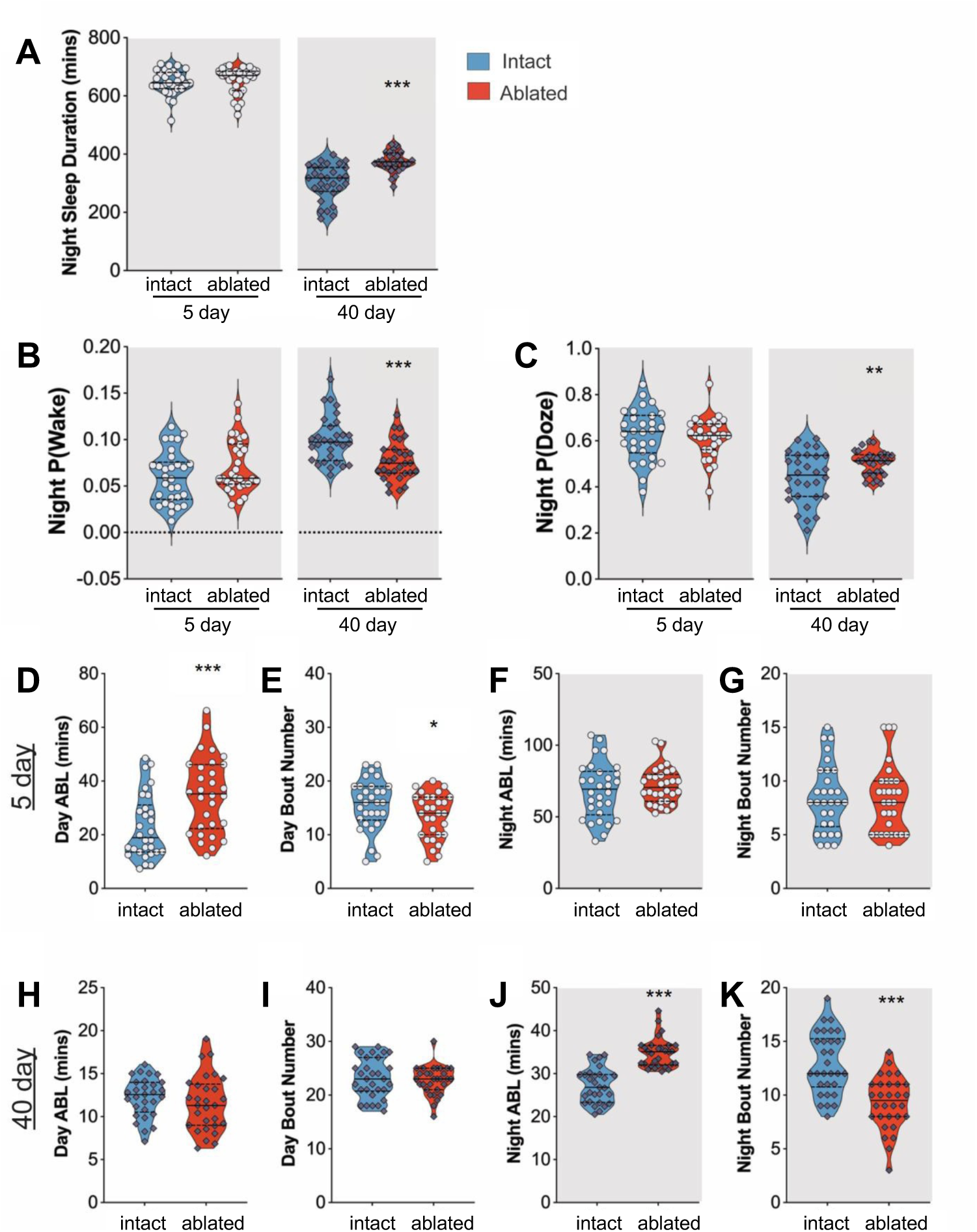
Sleep architectures differ from 5-day- and 40-day-old flies post neuronal injury. (**A**) Nighttime sleep duration was significantly increased in 40-day ablated flies following antennal axotomy (P=0.0002) but not in 5-day ablated flies (P=0.6031). (**B**) Nighttime wake probability, P(Wake), significantly decreased following ablation in 40-day-old flies (P=0.0003), but not 5-day-old flies (P=0.1171). **(C)** Nighttime sleep probability, (P)Doze, significantly increased following ablation in 40-day-old flies (P=0.0057) but not 5-day-old flies (P=0.6501). (**D**-**G**) Sleep architecture in 5-day-old flies post neuronal injury. Day average bout length (ABL, ZT2-ZT12 on the injury day) was significantly elevated (**D**, P=0.0007) but night ABL (ZT12-ZT0 on the injury day) was not significantly changed (**F**, P=0.4128). Day bout number (ZT2-ZT12 on the injury day) was declined (**E**, P=0.0369) but night bout number (ZT12-ZT0 on the injury day) was not significantly changed (**G**, P=0.8378). (**H**-**K**) Sleep architecture in 40-day-old flies post neuronal injury. Day ABL (ZT2-ZT12 on the injury day) was not significantly changed (**H**, P=0.3508) but night ABL was significantly elevated (**J**, P<0.0001). Day bout number (ZT2-ZT12 on the injury day) was not significantly changed (**I**, P=0.6056) but night bout number (ZT12-ZT0 on the injury day) was significantly increased (**K**, P<0.001). Gray boxes denote nighttime. *P<0.05; **P<0.01; ***P<0.001. Error bars indicate ± SEM.

**Supplementary Figure 3.**
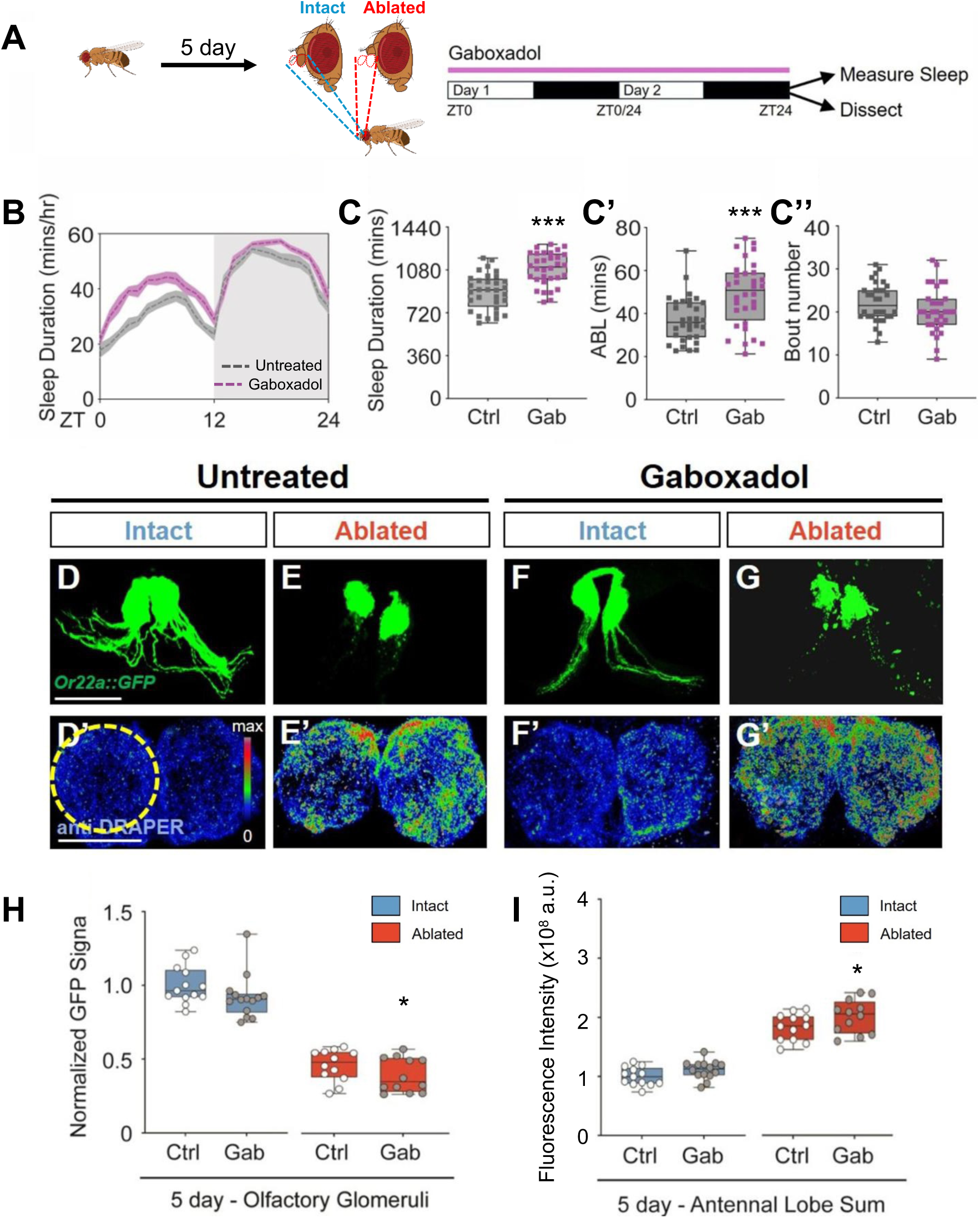
Gaboxadol treatment elevates glia-mediated clearance in 5-day-old flies. (**A**) Diagram of gaboxadol treatment and subsequent behavior monitoring and sample preparation. (**B**) Sleep profile of 5-day control and gaboxadol treated flies. (**C**) Total sleep and (**C’**) average bout length was significantly increased by gaboxadol treatment in 5-day-old flies (P<0.001 and P=0.0005, respectively), while no significant difference in (**C’’**) bout number (P=0.1652). (**D-G**) GFP signal labeled by Or22a::GFP and (**D’-G’**) immunostaining for DRAPER in the antennal lobes in gaboxadol treated and untreated intact or axotomized 5-day-old flies. Scale bar denotes 50 μm. (**H**) There was a significant reduction in GFP intensity in ablated 5-day-old gaboxadol treated flies compared to all other groups (P=0.0400). (**I**) Draper levels were significantly elevated in ablated 5-day-old gaboxadol treated flies compared to all other groups (P=0.0493). *P<0.05; **P<0.01; ***P<0.001. Error bars indicate ± SEM.

**Supplementary Figure 4.**
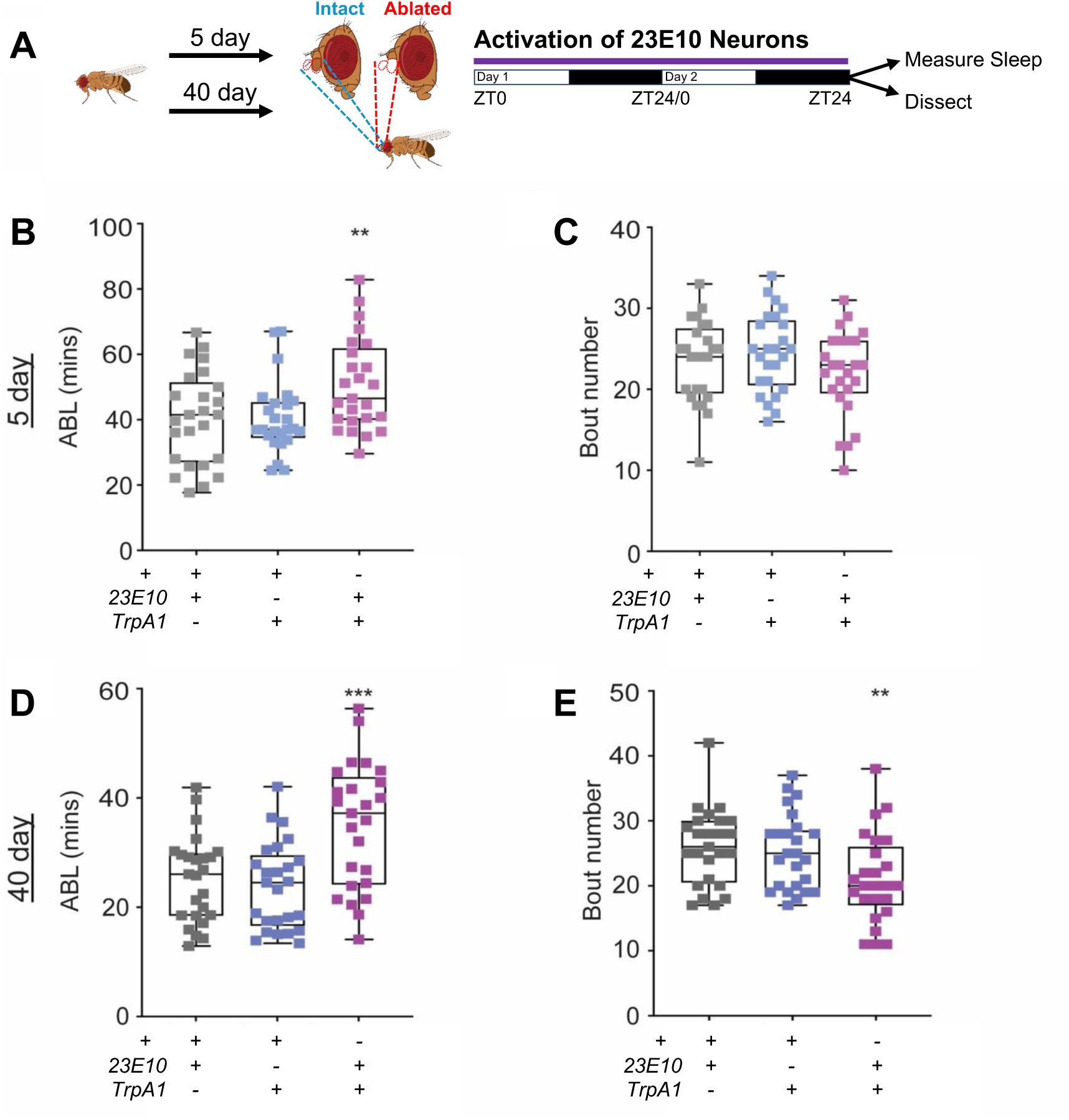
Sleep architectures in 5- and 40-day-old flies by genetic sleep induction. (**A**) Diagram of sleep induction by thermogenetic activation of 23E10 neurons and subsequent sleep measurement and quantification of glial engulfment. (**B**,**C**) Compared to controls, average bout length (ABL, **B**) significantly increased (P=0.0138) in 5-day-old 23E10>TrpA1 flies (P=0.0065), while (**C**) bout number did not significantly differ (P=0.2762). (**D**,**E**) Compared to controls, average bout length (ABL, **B**) significantly increased (P=0.0138) in 40-day-old 23E10>TrpA1 flies (P=0. 0002), while (**C**) bout number significantly decreased (P=0.0086). n=30-35. *P<0.05; **P<0.01; ***P<0.001. Error bars indicate ± SEM.

**Supplementary Figure 5.**
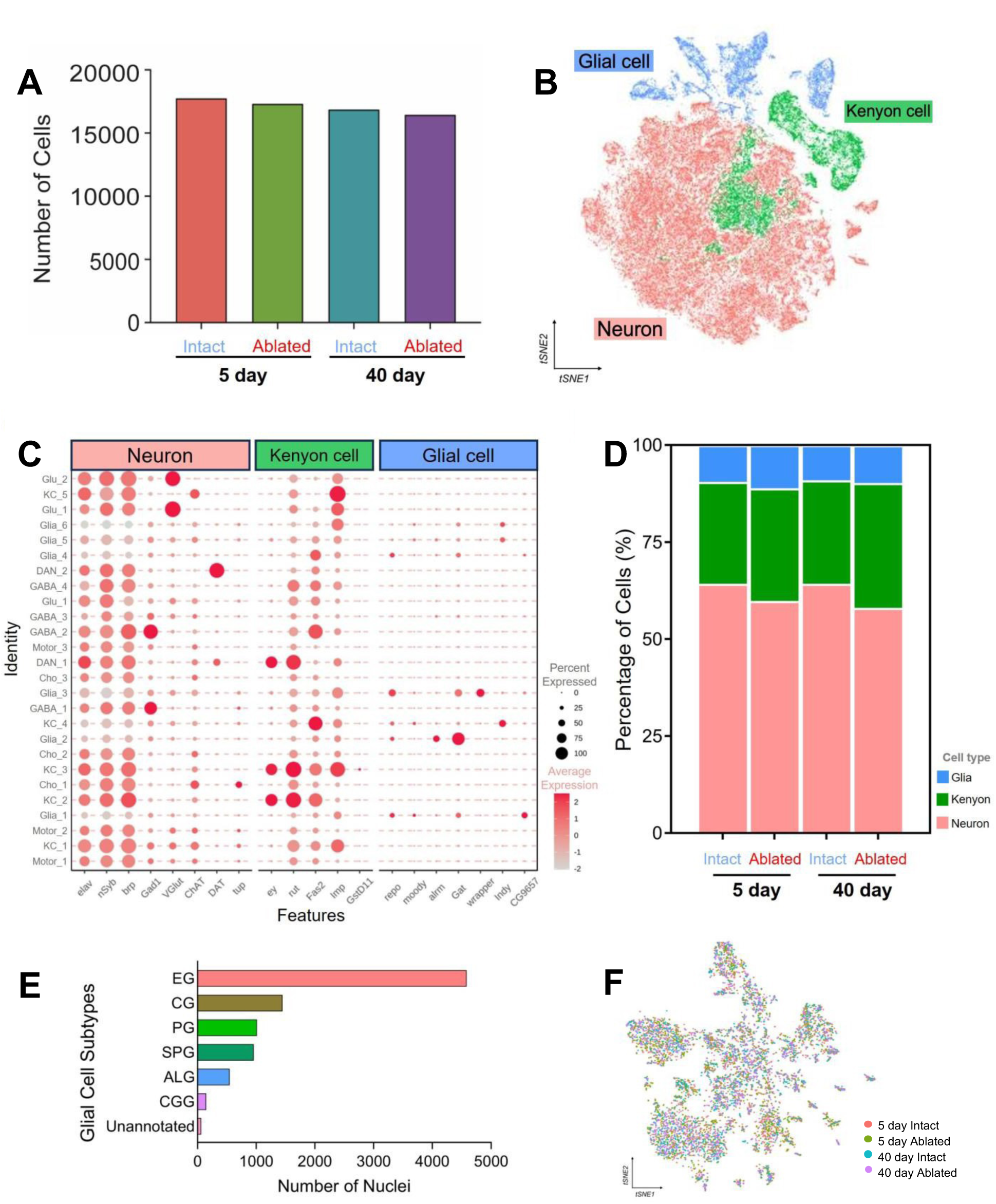
snRNA-sequencing of central brains in young and aged intact or ablated flies. (**A**) Number of nuclei collected from each pooled replicate. (**B**) t-SNE visualization of the 3 unique clusters: glial cells, Kenyon cells and neurons. (**C**) Maker genes expressed across the three cell types. (**D**) Percentage of cells for each of the three main groups for each sample. (**E**) Number of nuclei in each of seven glial cell subtypes. (**F**) t-SNE visualization of glial cells by four group samples.

**Supplementary Figure 6.**
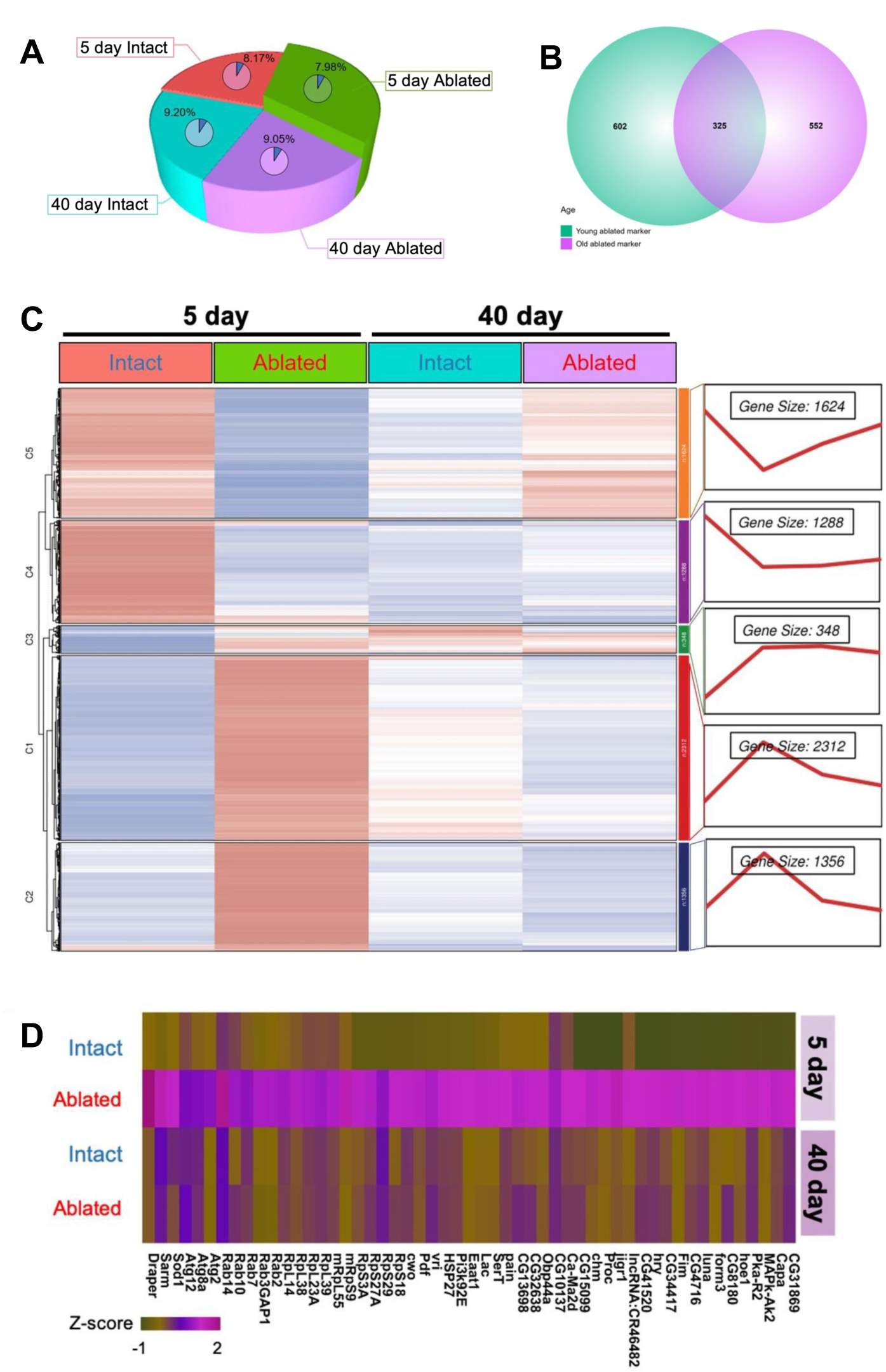
Differential gene expression clustering via pseudobulk analysis for olfactory ensheathing glia cells across 5- and 40-day-old intact and ablated flies. **(A)** The percentage of ensheathing glia across four sample groups and proportion of *Sarm* and *draper* co-expressed cells in each group. **(B)** Number of genes significantly upregulated in young or old ablated flies compared to intact controls. **(C)** Expression pattern clustering of all genes expressed in olfactory ensheathing glia via pseudobulk differential gene expression analysis. (**D**) Expression heatmap of selected candidate genes from the overlapping list of 602 unique 5-day ablation markers and genes in Cluster 2. (**E**-**H**) t-SNE visualization of selective marker genes for other glial subtypes on glia cohort, cortex glia (**E**, CG), perineural glia (**F**, PG), subperineural glia (**G**, SPG) and astrocyte-like glia (**H**, ALG). (**E’**-**H’**) *Sarm* and *draper* co-expressed cells highlighted on other glial subtypes, including CG (**E’**), PG (**F**’), SPG (**G**’) and ALG (**H**’).

## Notes

### Competing Interest Statement

The authors have declared no competing interest.

